# Cavitation Induced Fracture of Intact Brain Tissue

**DOI:** 10.1101/2022.03.15.484522

**Authors:** Carey E. Dougan, Zhaoqiang Song, Hongbo Fu, Alfred J. Crosby, Shengqiang Cai, Shelly R. Peyton

**Affiliations:** Chemical Engineering Department, University of Massachusetts, Amherst, USA; Mechanical and Aerospace Engineering Department, University of California, San Diego, USA; Polymer Science and Engineering Department, University of Massachusetts, Amherst, USA

**Keywords:** traumatic brain injury, fracture toughness, fracture energy, hydraulic fracture

## Abstract

Nonpenetrating traumatic brain injuries (TBI) are linked to cavitation. The structural organization of the brain makes it particularly susceptible to tears and fractures from these cavitation events, but limitations in existing characterization methods make it difficult to understand the relationship between fracture and cavitation in this tissue. More broadly, fracture energy is an important, yet often overlooked, mechanical property of all soft tissues. We combined needle-induced cavitation (NIC) with hydraulic fracture models to induce and quantify fracture in intact brains at precise locations. We report here the first measurements of the fracture energy of intact brain tissue that range from 1.5 to 8.9 J/m^2^, depending on the location in the brain and the model applied. We observed that fracture consistently occurs along interfaces between regions of brain tissue. These fractures along interfaces allow cavitation-related damage to propagate several millimeters away from the initial injury site. Quantifying the forces necessary to fracture brain and other soft tissues is critical for understanding how impact and blast waves damage tissue *in vivo* and has implications for the design of protective gear and tissue engineering.

**Significance:** Mild injuries associated with concussion and blast waves cause tearing of brain tissue, which leads to traumatic brain injury (TBI). TBI is a leading cause of death and disability among children and young adults in the U.S., with 1.5 million Americans reporting a TBI each year. We introduce a novel approach to visualize these tears in intact brain tissue, and report the energies associated with brain fracture. Quantifying the fracture energy of brain, as we have done here, is critical to understand the forces from injury that lead to TBI.

## 1. Introduction

According to the CDC, there were 223,050 traumatic brain injury (TBI)-related hospitalizations in 2018, and 60,611 TBI-related deaths in 2019 [1], [2]. For military members, explosive blasts produce the majority (66%) of injuries resulting in TBI [3]. Characterizing the mechanical properties of brain is essential to understand how external forces such as blast waves result in TBIs. Unlike other tissues in the body, brain lacks the loadbearing fibrillar proteins of the extracellular matrix (ECM), like collagen, making it ultra-soft with a modulus ~1 kPa [4], [5]. The modulus varies between grey and white matter, and the corona radiata is stiffer than the cortex, thalamus, and corpus callosum [6], [7]. Within specific regions, the modulus changes over the course of an organism’s lifetime [8]. Under loading, brain shows both a directional and strain rate dependency [9], likely due to its porosity [10] and heterogeneity such as: interfaces between the ECM and blood vessels, and along boundaries between different regions (*e.g*., between white and gray brain matter).

Despite this extensive mechanical characterization of brain tissue, we know little about injury propagation. The delicate nature of brain tissue makes it susceptible to tearing when exposed to external forces, yet few studies have quantified the precise forces required to tear or fracture brain [11]. Understanding fracture mechanics of brain tissue is crucial to characterize the extent of injury and potentially link cavitation related mild TBI with neurodegeneration.

Fracture energy measures the energy required to open a crack in a material and is closely, or often directly, related to the material’s fracture toughness [12]. Knowing the fracture energy of brain is required to relate impact and blast wave forces with tissue damage and TBI. During blast, waves travel through the head and reflect off the skull, causing regions of high and low pressure that result in bubble formation and collapse, *aka* cavitation. Recent studies show that bubble growth can lead to fracture if the driving force reaches a critical value [13]. Fracture has been observed in laboratory blast wave experiments, where tearing of rat brain tissue along interfaces resulted in permanent realignment of tissue [14]. To our knowledge there has been no controlled method to quantitatively link cavitation pressures to tissue fracture energies in intact brains. Achieving this required us to combine microscale, precise cavitation of brain and application of hydraulic fracture models. We report here for the first time the fracture energy of intact brain tissue, which determines the extent of TBIs.

## 2. Materials and Methods

### Animals

A total of 57 male and female BALB/c, a mix of homozygous (−/−) or heterozygous (+/−) nude mice (6-8 weeks old; The Jackson Laboratory, Bar Harbor, ME USA) were used. The mice were housed in a pathogen-free environment with a 12:12-hour light:dark cycle and controlled humidity and temperature, with unlimited access to food and water. We first sacrificed mice, excised their brains 4-10 at a time, and stored the tissues in ice cold 1X Hank’s balanced salt solution (Gibco, Waltham, MA USA) prior to NIC. All protocols applied in the experiments were approved by the Institute of Animal Care and Utilization Committee at University of Massachusetts Amherst. Fresh porcine lung (Animal Technologies, Tyler, TX USA) arrived on ice and was stored hydrated on ice until testing was performed.

### Synthesis of Alginate Gels

Alginic acid sodium salt from brown algae (Sigma-Aldrich, St. Louis, MO USA) was dissolved in distilled water (97% vol/vol water/alginate for the hard gels and 98.5% vol/vol water/alginate for the soft gels) for 12-24 hours on a stirrer. A 0.75 M calcium sulfate dihydrate (CaSO4) (Sigma-Aldrich) in water solution was stirred continuously. 10 wt% calcium sulfate solution was injected into 20 mL of alginate solution. Two 20 mL syringes (Thermo Fisher Scientific, Waltham, MA USA) were attached with a syringe connector (C-U Innovations, Chicago, IL USA) and solution was rapidly dispensed between two the syringes 6 times to mix before injection into scintillation vials (Thermo Fisher Scientific) or 3D printed (Taz 4, Lulzbot. Loveland, CO USA and Ender-3 Pro, Creality, Shenzhen China) pure shear molds (10 cm x 2 cm x 4 mm). Alginate solutions were covered and allowed to fully polymerize at room temperature overnight.

### Needle-Induced Cavitation (NIC)

The collected mouse brains were stored in ice cold 1X Hank’s balanced salt solution (Gibco) until NIC, and all mouse brain experiments were conducted within 30 minutes post-harvesting. For the NIC experiment, the cavitation fluid was distilled water with 1:1000 vol/vol DAPI (Sigma-Aldrich) and 1:100 vol/vol 200 nm far red carboxylate functionalized latex beads (Thermo Fisher Scientific). For the alginate NIC experiment, the cavitation fluid was distilled water with food coloring (McCormick’s, Baltimore, MD USA) for visualization. Just prior to testing, lung tissue was cut into 1-inch squares using surgical scissors (Thermo Fisher Scientific) and placed in a custom 3D printed (uPrintSE+, AET Labs, Essex, MA USA) specimen holder. For the lung experiments, the cavitation fluid was distilled water. For all materials, control of the pressure was achieved using a 1 mL glass syringe (Hamilton, Reno, NV USA) in a Nexus 6000 syringe pump (Chemyx, Stafford, TX USA). During NIC experiments, the real time pressure was monitored using a Px409-015 GUSBH (Omega, Norwalk, CT USA) and interfaced with a custom LabView program. The depth of needle insertion was controlled using an actuator (Texture Technologies, Hamilton, MA USA). Each NIC experiment took approximately 1 minute. After NIC, brains were immediately placed in 10% formalin (Thermo Fisher Scientific) and stored in a 4°C refrigerator for 48-72 hours prior to slicing.

### Fracture Energy Calculations

From the NIC experiment, we can use the critical pressure *P_c_* to calculate the modulus *E* from the equation [15]

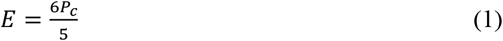

We ignored the effects of surface tension between water and tissue. We calculated the plane strain modulus *E*′ from the equation [16]:

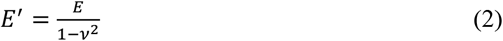

where *v* is Poisson’s ratio and approximately equal to 0.5 for brain tissue [17], 0.4 for lung tissue [18], and 0.23 for alginate gels [19] for conditions consistent with our measurement approach. From the raw pressure-time data obtained through NIC, we plotted the data from the critical pressure to when the pump was turned off (Figure 1a). We set the critical pressure time to be zero and plotted log(P) versus log(t) to determine the slopes (Figure 1b). In cases of obvious slope changes, we used the second, steeper slope, as this data represents the stable fracture propagation regime. Based on the slope of the log(P) versus log(t) plot, we applied either the plane strain (slope ~ - 1/3) or axisymmetric (slope ~ −1/5) model. The pressure-time relation for the plane strain case is

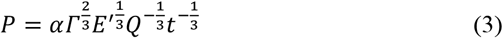

where *α* is a constant 0.587, *P* is pressure, *Q* is flowrate, *Γ* is fracture energy, and *t* is time [16]. Solving for fracture energy, we have the following equation

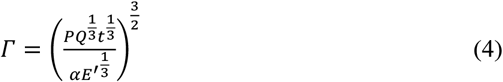

**Figure 1.**
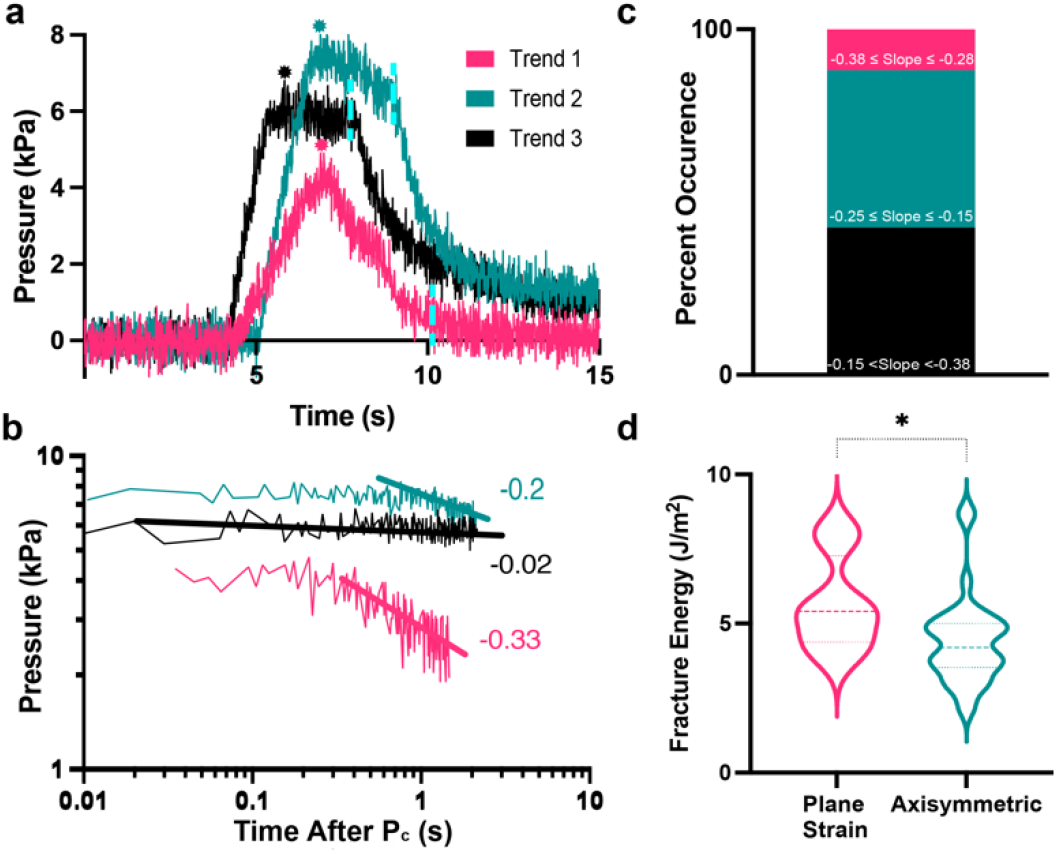
Intact brain fracture energy using NIC. **a**. Three observed pressure-time trends (1: pink; 2: teal, and 3: black) in experimental data have different slopes after a maximum pressure is reached. Burst shape indicates the timepoint for the critical event and blue dashed line indicates when the syringe pump is turned off. **b**. Log-log data transformation from 1a between the maximum pressure value to when the pump was turned off. Trendlines show slopes of typical trends observed. c. The distribution of experiments with slopes in the ranges: −0.38 ≤ slope ≤ −0.26(pink), −0.25 ≤ slope ≤ −0.15 (teal), or −0.15 < slope < −0.38 (black). **d**. The fracture energies of mouse brain calculated using the plane strain or axisymmetric hydraulic fracture models.

The pressure-time relation for the axisymmetric model is

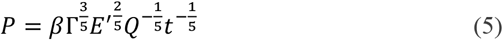

where *β* is a constant 1.41 [16]. Thus, the fracture energy is:

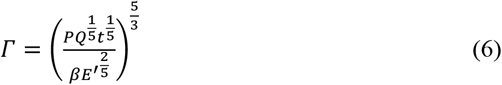

We used a least-squares method to fit a line to the data from the critical pressure to when the pump was turned off. We used the y-intercept pressure P(log(t=0)) to calculate the fracture energy *Γ* by using Eqs. (4) and (6).

### Brain Slicing and Imaging

After NIC and formalin fixation, mouse brains were submerged in 15% sucrose (Thermo Fisher Scientific) in 1X PBS for 6-12 hours and then in 30% sucrose in 1X PBS for 24 hours. Brains were immersed in HistoPrep OCT (Thermo Fisher Scientific) for 10 minutes before flash freezing and stored in the −80°C freezer until ready for slicing. Tissue slicing was performed using a Leica CM1860 cryostat (Leica, Wetzlar, Hesse Germany) and slices were arranged on Superfrost Plus Microscope Slides (Thermo Fisher Scientific), covered with gelvatol (Supplemental material) aqueous medium and a #1.5 glass coverslip (Corning, Corning, NY USA), and sealed with clear fingernail polish (Sally Hansen, Morris Plains, NJ USA). Imaging was performed on a Spinning Disk Axio Observer.Z1 (Zeiss, Oberkochen, Germany).

### Alginate Gel Pure Shear

We used a traditional method, pure shear, to validate NIC measured fracture energy of alginate gels. The pure shear tests were conducted on rectangular alginate gels by applying uniaxial tensile loading along the height direction of notched and unnotched samples [20]. Transparency paper (Staples, Framingham, MA USA) was super glued (Gorilla, Cincinnati, OH USA) to both sides of the hard alginate gel at the top and bottom and clamped into custom fixtures. The soft alginate gels were directly clamped at the top and bottom. Tensile loading was performed using an actuator (Texture Technologies) and both notched and unnotched gels were stretched at a rate of 100 μm/s while force was measured with a 50 N load cell in real time.

### Numerical Simulations

We developed a finite-element model (FEM) to simulate the hydraulic fracture of brain tissues by using Abaqus (version 2017. Dassault System, Johnston, RI USA). The plane strain and axisymmetric models were set up in ABAQUS/Standard with a size of 100 mm by 100 mm. We used an initial crack size with a length of 0.1 mm with extra fine mesh near (< 10% of the sample size) the initial crack and coarse mesh far (> 10% of the sample size) from the crack. For the plane strain model, brain tissue was modeled as a linear poroelastic material with a void ratio of 0.25, defined as the ratio between volume of the empty space and solid, using the pore pressure plane strain elements (CPE4P: 4-node bilinear displacement and pore pressure). The crack growth in the material was modeled by a cohesive zone model using a layer of cohesive element COH2D4P (6-node displacement and pore pressure twodimensional cohesive element) with zero thickness. For the axisymmetric model, brain tissue was modeled as a linear poroelastic material with a void ratio of 0.25 by using the cohesive element CAX4P (4-node bilinear displacement and pore pressure), and the interaction between hydraulic fracture and natural fracture was simulated by cohesive zone model with cohesive element COHAX4P (6-node displacement and pore pressure axisymmetric). We used a modulus of brain tissue of 20 kPa, the interfacial strength ranging from 0.3 to 2 kPa, and the fracture energy ranging from 0.0625 to 1 J/m^2^ for our simulations. We chose values that slightly deviate from the experimental values, because FEM simulations do not converge if we use the exact parameters for the brain.

### Statistical Analysis

Unpaired Mann-Whitney t-tests were performed with GraphPad Prism (GraphPad Software, La Jolla, CA USA) on all data unless otherwise noted. Ordinary one-way ANOVA tests were performed using Prism (GraphPad Software) on Supplemental Figure 2a-b, Supplemental Figure 3a, and Supplemental Figure 6. *, **, ***, and **** indicate statistically significant differences with p ≤ 0.05, p ≤ 0.01, p ≤ 0.001, and p ≤ 0.0001, respectively.

## 3. Results

### Fracture Energy of Murine Brain

We started recording NIC data prior to pressurization. A critical event occurred at a maximum pressure, followed by either a decrease or levelling off before the syringe pump was turned off (Figure 1a).

We used data from the NIC experiments to calculate the modulus of brain tissue. We measured the modulus of different regions of brain tissue using NIC as previously described [15]: cortex 4.5 ± 1.5 kPa, thalamus 7 ± 1.5 kPa, hypothalamus 7.9 ± 1 kPa, and cerebellum 5 ± 1 kPa (Supplemental Figure 1a). These moduli obtained via NIC are higher than reported using other techniques, a trend also observed by others [5]. In our NIC modulus calculations (E=6P_c_/5), we ignored the effects of surface tension, and assumed the samples were neo-Hookean solids.

Analysis of these pressure-time curves revealed three typical trends after a critical pressure: a sharp drop (pink, Trend 1), gradual drop (teal, Trend 2), or constant pressure (black, Trend 3, Figure 1a). Based on the log-log transformed slopes of these raw data between the critical pressure and when we turned the pump off, we applied one of two different hydraulic fracture models: plane strain or axisymmetric. The asymptotic solution for the plane strain model has a slope ~ −1/3 and it is ~ −1/5 for the axisymmetric model (Figure 1b, Eqs. (1) and (3)). Fracture energies were calculated based on ranges of slopes: between −0.28 and −0.38 for the plane strain model and between −0.15 and −0.25 for the axisymmetric model (Figure 1c and Supplemental Figure 2). Among 68 experiments on mouse brain, the axisymmetric model applied to 30 cases and plane strain applied to 8 cases, and roughly 40% of experiments fit neither model. Using the asymptotic solutions for both the plain strain and axisymmetric model, we calculated the average fracture energy of murine brain to be 5.6 ± 1.6 J/m^2^ for those cases that fit the plane strain model and 4.4 ± 1.5 J/m^2^ for axisymmetric cases (Figure 1d). Fracture energy measurements were consistent across mouse sex and genotype (Supplemental Figure 3).

We expected that the fracture energies would correlate with the modulus of the different brain regions. Despite statistically significant differences in moduli between brain regions measured using NIC, the fracture energies were consistent (Supplemental Figure 1b). We were surprised that regardless of the model applied, the fracture energies were similar. Therefore, we performed a test, applying the “incorrect” model to each data set. The fracture energy calculated using the axisymmetric model for data that fits the plane strain model was slightly lower than when calculated using the correct model, but the fracture energy calculated using the plane strain model for data that fits the axisymmetric model was significantly higher than when calculated using the correct model (Supplemental Figure 4). Both the plane strain and axisymmetric are simplified models, so it could be expected that they would result in similar values.

### Visualizing Fracture in Mouse Brain Along Tissue Interfaces

To visualize these fracture events in brain during NIC, we included a DAPI nuclear cell dye (blue) and fluorescent beads (red) in the injection fluid (Figure 2). We imaged several consecutive slices of tissue and observed that the fracture paths extended far (up to 7 mm) from the initial cavitation site (beginning at 1.8mm depth). Fracture consistently occurred along the interface of the hippocampus region (white arrow). This is significant given that injuries to the hippocampus can result in memory loss, depression, and epilepsy [21]. This finding also supports broad research suggesting that the hippocampal region is disproportionately affected by TBI [22]. Through visual confirmation, we conclude that fracture is the primary mode of damage through brain tissue after cavitation.

**Figure 2.**
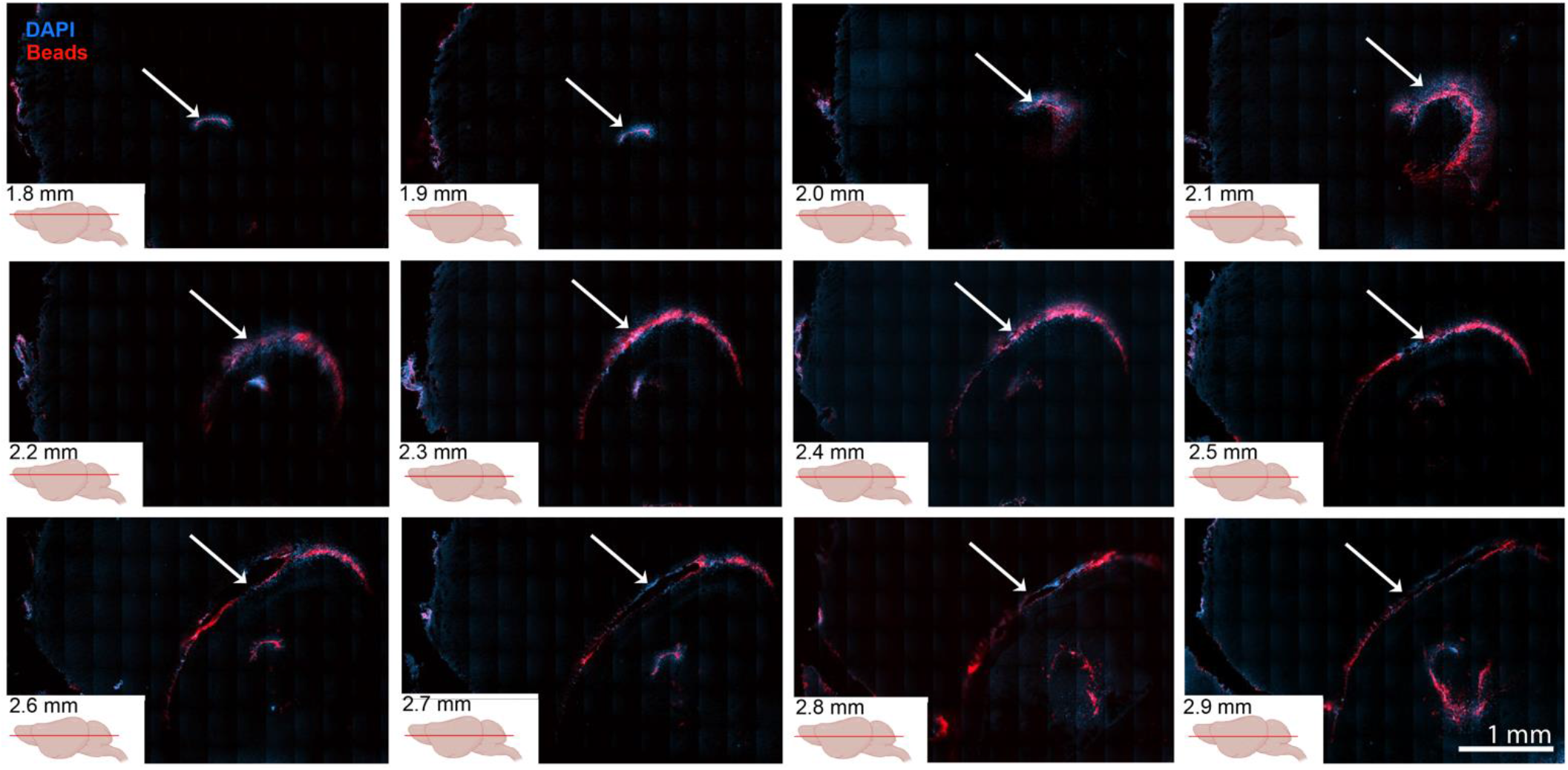
Fracture occurs along tissue interface. Through subsequent 100 μm thick horizontal slices of mouse brain, starting in the top left image (1.8 mm deep) and ending with the bottom right image (2.9 mm deep), fracture occurs along the hippocampus interface (white arrows) in the brain visible by both DAPI staining (blue) and fluorescent beads (red). The scale bar is 1 mm.

### Validation of NIC-Induced Fracture with a Model Material

To the best of our knowledge, the fracture energy of intact brain tissue has not been previously reported. Therefore, we validated our approach using a model alginate gel, which allowed us to measure and visualize fracture in real time [23]. We prepared two formulations of alginate gels (3 and 2.5 vol%) for NIC and pure shear testing, which resulted in moduli of 17.8 ± 7.6 (hard) and 2.9 ± 0.8 kPa (soft) respectively (Figure 3a). We directly observed NIC- and pure shear-induced fracture for both formulations during testing (Supplemental Videos 1–4). The fracture energies were consistent between NIC and pure shear (Figure 3b-c): 0.3 ± 0.1 J/m^2^ (NIC) and 0.2 ± 0.2 J/m^2^ (pure shear) for soft alginate gels, and 4.3 ± 1.7 J/m^2^ (NIC) and 4.2 ± 1.0 J/m^2^ (pure shear) for hard gels. Given the consistency of fracture energies between NIC and a traditional pure shear technique in these cases where we can directly visualize fracture, we are confident that NIC is a valid method to measure the fracture energy of brain tissue where real-time visualization of fracture is not possible.

**Figure 3.**
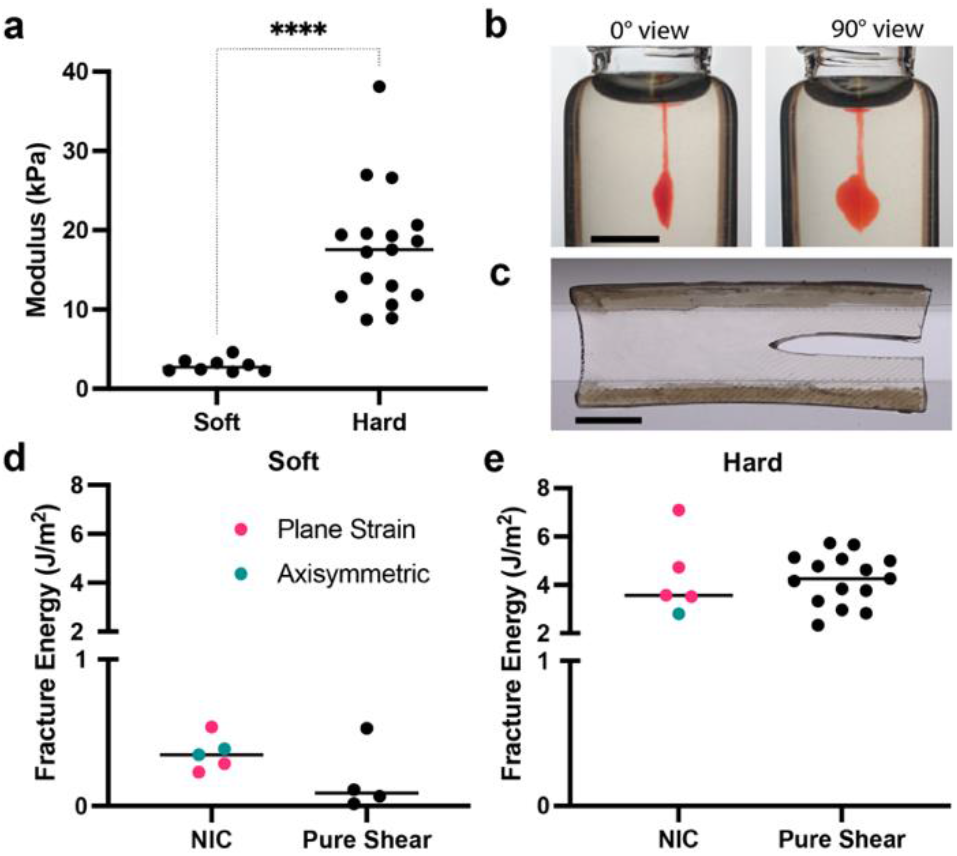
Alginate gel fracture energies in NIC consistent with pure shear. **a**. NIC measured moduli for soft and hard alginate gels. **b**. Front and side views of NIC induced fracture in a hard alginate gel. **c**. Picture of pure shear testing of a hard alginate gel. Fracture energy of the **d**. soft and **e.**hard alginate gels measured by NIC and pure shear testing. For NIC experiments the values were calculated using either the plane strain (pink) or axisymmetric (teal) model. Scale bars represent 2 cm.

We measured the fracture energy of porcine lung tissue to confirm that we could apply this approach to another complex tissue. Using NIC, we measured the modulus of porcine lung to be 8.31 ± 3.72 kPa, which agrees with other reports [24], and we calculated the average fracture energy of porcine lung (N=25) 13.8 ± 7.3 J/m^2^ via the plane-strain model (N=8) and 7.6 ± 5.2 J/m^2^ from the axisymmetric model (N=5) (Supplemental Figure 5a-b). Although lung tissue has been studied in tension at strains that result in fracture, this is the first reported value for a fracture energy of lung [25].

### Validation of Hydraulic Fracture Model by Finite Element Simulation

Analytical solutions for the toughness-dominated hydraulic fracture of rock with plane strain or axisymmetric assumptions were given in Eqs. (3) and (5) [16]. We used the cohesive traction-separation law to study fracture in brain tissues (Figure 4a, Supplemental Materials). To derive the analytical solution, the material was assumed to be infinitely large, and the crack tip was assumed to be sharp [16]. However, in experimental hydraulic fracture of brain (Figure 4b), the sample size was finite, and the crack tip could be blunted. Therefore, we conducted finite element simulations to examine the validity of the analytical solutions using a finite sample size and a blunted crack tip (Figure 4c and f). We calculate the fractocohesive length *r_c_*

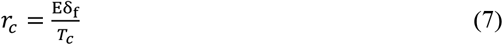

where *δ_f_* is the separation at failure and *T_c_* is the interfacial strength (Figure 4a). When *r_c_* was much smaller than the crack length (sharp crack tip), the slope of the pressure-time curve in a log-log plot approached −1/3 for plane strain and −1/5 for axisymmetric. Otherwise, the slope was larger than −1/3 (plane strain, Figure 4d-e) or −1/5 (axisymmetric, Figure 4g-h). We found that calculated fracture energies based on either model were within 50-200% of the fracture energies simulated with input values similar to experimental values (Supplemental Figure 5). Thus, Eqs. (4) and (6) provide a good estimation of fracture energy for our NIC experiments.

**Figure 4.**
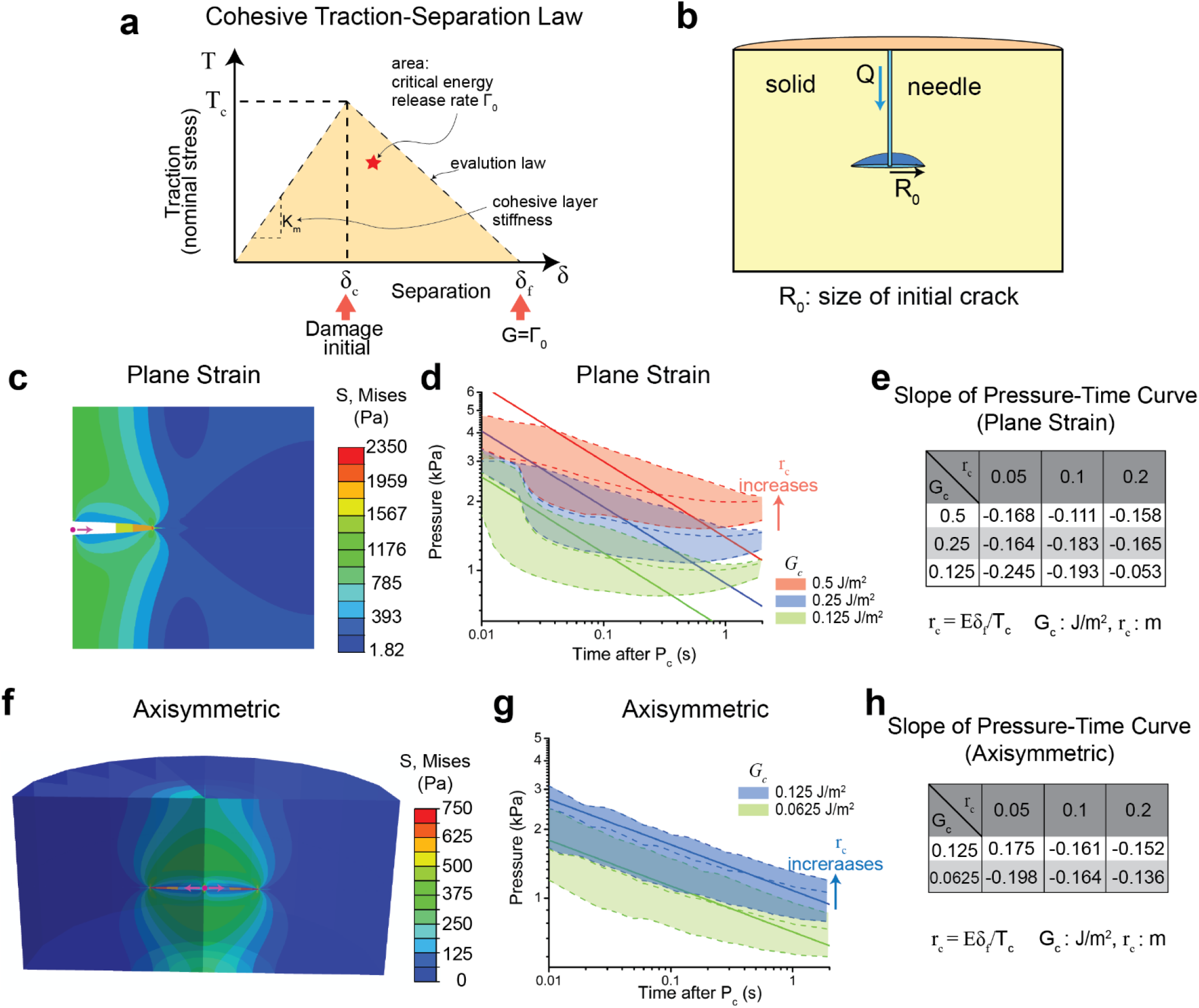
Numerical simulations of hydraulic fracture in brain tissue. **a**. Cohesive traction-separation law is applied to study fracture in brain tissue. **b**. A schematic of NIC-induced fracture. **c**. Stress distribution of finite element simulations for the plane strain case. **d**. The pressure-time curves for varying material properties for the plane strain case. **e.**The slope of pressure-time curves with varied material properties for the plane strain case. **f.**Stress distribution of finite element simulations for the axisymmetric case. **g**. The pressure-time curves for varying material properties for the axisymmetric case. **h**. The slope of pressuretime curves with varied material properties for the axisymmetric case. Dashed lines in **d** and **g** are the results of FE simulations, and solid lines are the results from theoretical analysis.

## 4. Discussion

Fracture energy values have been reported for skin, cartilage, muscle, liver, and vocal fold tissue [26]–[29]. The fracture energy of brain tissue, however, has only been measured from dissected pieces [30], and not for intact brain. Traditional techniques for measuring the fracture energy of soft tissues, such as pure shear [20], single-edge crack [31], or tearing tests [32], are challenging for many reasons [33]. One limitation is the tissue sample size and availability for bulk mechanical measurements. Another challenge is achieving the specific geometric requirements necessary for using these more conventional methods. Cutting or shaping the tissue to fit such geometric constraints also interrupts the structural integrity of tissues. Lastly, it is difficult to avoid damage to tissue during transport and handling required for traditional mechanical testing approaches.

NIC is a technique that overcomes these challenges and allows us to induce cavitation and measure the fracture energy of intact tissues. The morphology of the voids initiated by NIC are spherical for elastic, homogeneous, and isotropic materials; however, depending on the loading mode and initial crack size, a fracture event may grow during the loading process [13], [34] making NIC well-suited to measure the localized fracture properties of brain tissue [13]. Many researchers have used NIC to measure the modulus of soft materials and biological tissues [15], [34], [43]–[52], [35], [53]–[56], [36]–[42], and some have adopted NIC techniques to characterize stiff materials and to understand the effects of viscoelasticity and loading conditions [54], [57], [58]. More recently, NIC has been recognized as a method to measure damage associated with cavitation events, which have implications for injuries related to impact or blast wave exposure [59]. Here we show that the modulus does not directly correlate with the fracture energies of those same materials (Supplemental Figure 7).

The pressure-time profiles during NIC of brain tissue resembled that of hydraulic fracture of rock, with one critical difference. Hydraulic fracture of shale rock is dominated by the fluid viscosity, namely, the external work overcomes the energy dissipation by the fluid flow [60]. In contrast, for brain tissue, fracture is dominated by tissue tearing. Whether the process is viscosity-dominated or fracture energy-dominated is determined by the dimensionless parameter *χ*

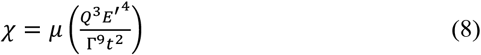

where *μ* is the viscosity of fluid, *Q* is the flowrate, *E* is the modulus, *Γ* is the fracture energy, and *t* is the injection time. In NIC of brain, *χ* « 1, and thus the hydraulic fracture in brain is fracture toughness dominated, enabling NIC to report the fracture energy.

We applied plane strain and axisymmetric hydraulic fracture models to our NIC experimental data to estimate the fracture energy of brain. This approach required injecting an incompressible Newtonian fluid at a constant rate to drive fracture in a permeability-ignored, infinite, brittle, and poroelastic solid. These assumptions are appropriate given the delicate nature of brain and the poroelastic behavior of brain parenchyma [17]. The fracture energy values (4-5 J/m^2^) we report for brain tissue are consistent across both models and reasonable given that brain is fragile. In comparison, the fracture energy of cartilage is ~1000 J/m^2^ [61]. Nearly 40% of our NIC experimental data fit with neither model. In linear elastic fracture mechanics, the crack tip is sharp, but because brain is viscoelastic, the crack tip may be blunted resulting in the pressure leveling off, in which case neither plane strain nor axisymmetric hydraulic fracture models can be used to calculate a fracture energy.

There is another important assumption in the hydraulic fracture models used: leak-off (loss of fluid into solid material) has a negligible effect. For brain tissue, the timescale of water diffusion through poroelastic brain can be estimated as

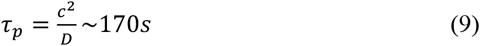

where the crack size *c* is approximately 1 mm, and the diffusion coefficient *D* of water in brain tissue is 3.2 × 10^-5^*cm*^2^/*s* [62]. The timescale of poroelasticity is much larger than the duration time of the injection, thus, the leakage of water was ignored. Strain rates associated with impact and blast waves are 36 to 241 s^-1^ [63]. It has been shown that strain rates in the range of 22 - 75 s^-1^ result in damage on a cellular and sub-cellular level [64] [65]. The time scales associated with fracture of brain tissue here were ~2 seconds, and resulted in fractures that measured up to 7 mm, a crack extension speed of maximally 3 mm/s. We expect that the fracture properties of brain tissue, at the time and length scales associated with NIC, will add to a growing body of research quantifying the mechanical properties of brain under varying loading conditions [5], [17], [66]–[68].

Both numerical simulations and experiments indicate that cavitation may be a damage mechanism contributing to TBI from blast [69]–[72]. The specific mechanism by which cavitation damages tissue is still unclear, be it bubble collapse, cell membrane interruption, blood-brain barrier disruption, or water jet formation [73]. Although cavitation-induced damage has been measured in brain slices and cell culture systems [74][76], there is still a missing link that connects cavitation to fracture. Our study provides this link, using cavitation to initiate fracture in murine brain to report on fracture energies relevant to mild TBI. With precise needle control, we capture localized mechanical and fracture properties with spatial variance. There is still a need to compare the *in vivo* fracture properties of brain with these *ex vivo* characteristics, and NIC is uniquely suited to measure *in vivo* tissue properties.

## Conclusions

We use needle-induced cavitation (NIC) in combination with hydraulic fracture models to gain insight into the local fracture mechanics and the propagation of cavitation damage in intact brain tissue. Cavitation-induced fracture in brain tissue extends for several millimeters beyond the initiation site along tissue interfaces. The average fracture energy of brain tissue is 4.7 ± 1.6 J/m^2^, with agreement between the two hydraulic fracture models we applied (5.6 ± 1.6 J/m^2^ for plane strain and 4.4 ± 1.5 J/m^2^ for axisymmetric). This is the first report of fracture energy of intact brain tissue. We conclude that both hydraulic fracture models are robust and provide good approximations for fracture energies in these complex tissue fracture events. Thus, combining NIC with hydraulic fracture models is ideal for calculating fracture energies of brain.

## Author Contributions

C.E.D and Z.S. contributed to drafting the manuscript, data collection, and data interpretation. H.F., A.J.C., and S.C. contributed to data analysis and interpretation, and editing the manuscript. S.R.P. contributed to conceptual design, data interpretation, and drafting the manuscript.

## Acknowledgements

This research was supported by a grant from Office of Naval Research (N00014-16-1-2872) to S.R.P, S.C., and A.J.C. C.E.D. was supported by an NIH funded Chemistry Biology Interface Fellowship (T32 GM008515 and T32 GM139789), and a Spaulding-Smith Fellowship from the UMass Amherst Graduate School. S.R.P. was supported by an Armstrong Professorship from UMass Amherst.

## Supplemental Material

**Supplemental Figure 1.**
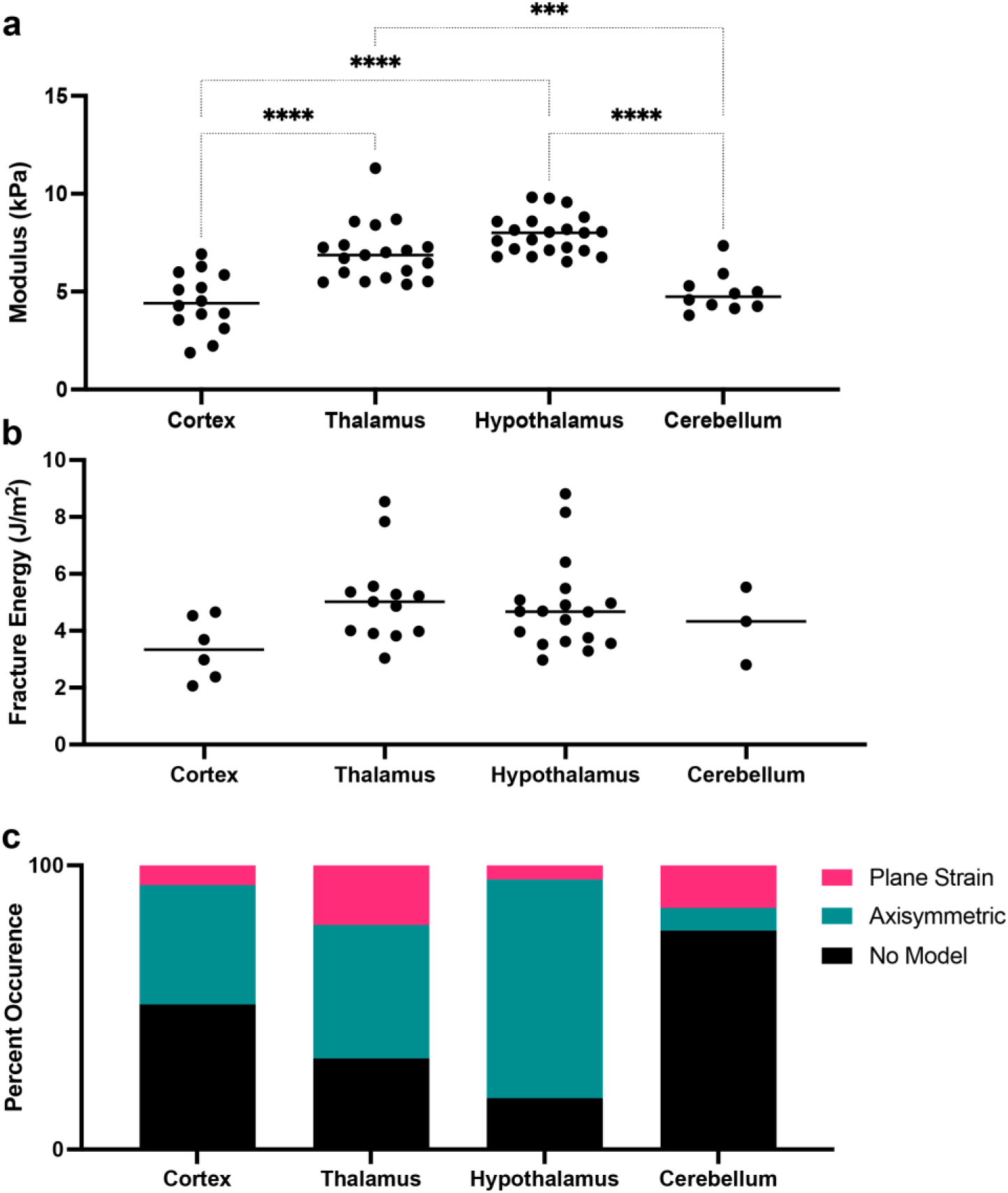
Fracture energy consistent across brain regions. **a.** The modulus measured using NIC varies between region, but the fracture energies **(b)** are consistent across different regions of the brain **c.** The distribution of the models applied for different regions of the brain.

**Supplemental Figure 2.**
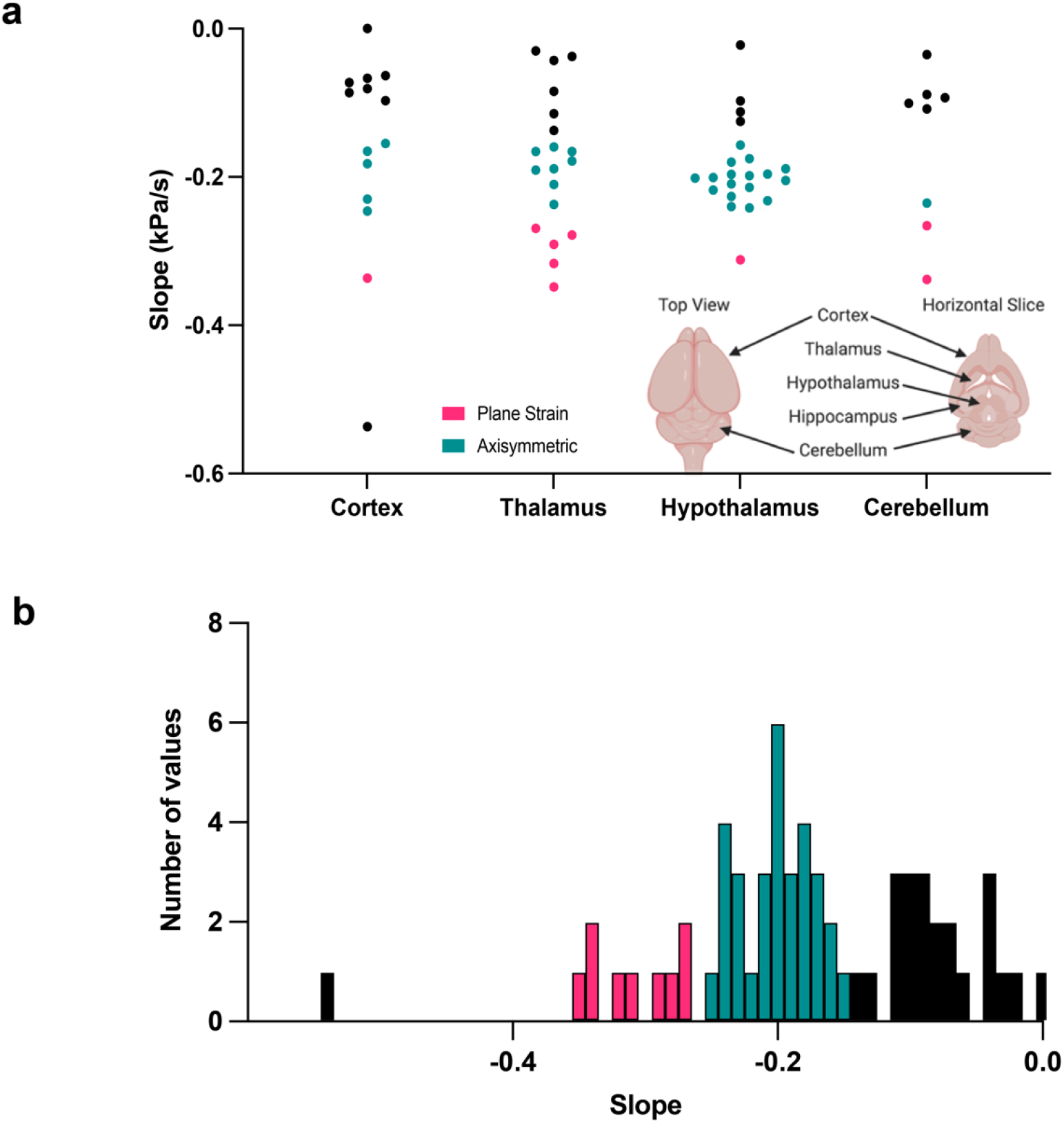
Slope distributions for log-log transformation of data. **a.** Range of slope values from the critical pressure to when the pump was turned off for all mouse brain NIC data for both plane strain (pink) and axisymmetric (teal) models across different regions of tissue (n=68). **b.** Frequency distribution of slopes for all NIC mouse brain experiments.

**Supplemental Figure 3.**
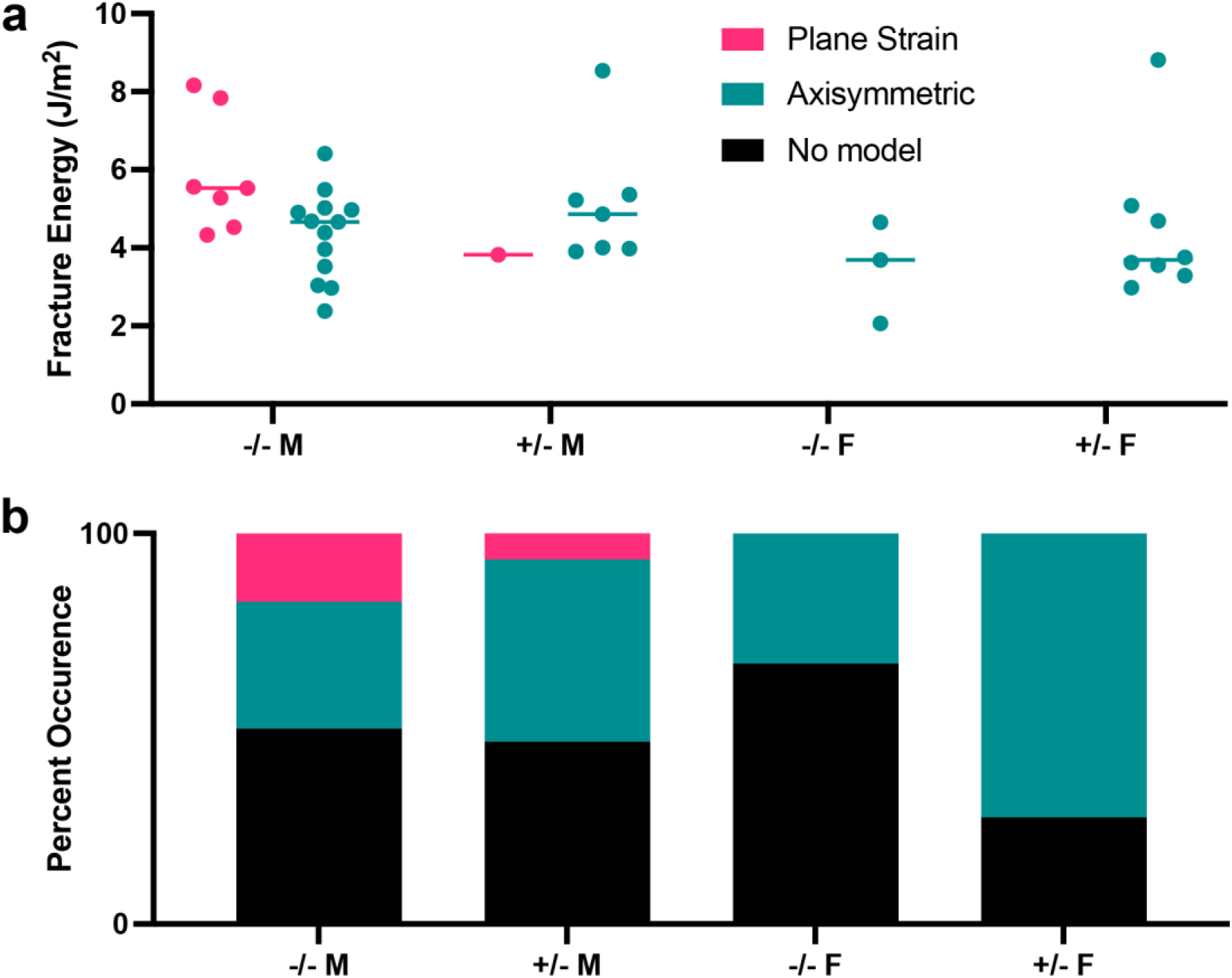
Fracture energy by sex/genotype of mouse. **a.** Neither genotype: homozygous (−/−) or heterozygous (+/−) nor sex: male (M) or female (F) change the fracture energy values for mouse brain. **b.** The distribution of models applied to experimental data based on sex and genotype of mouse.

**Supplemental Figure 4.**
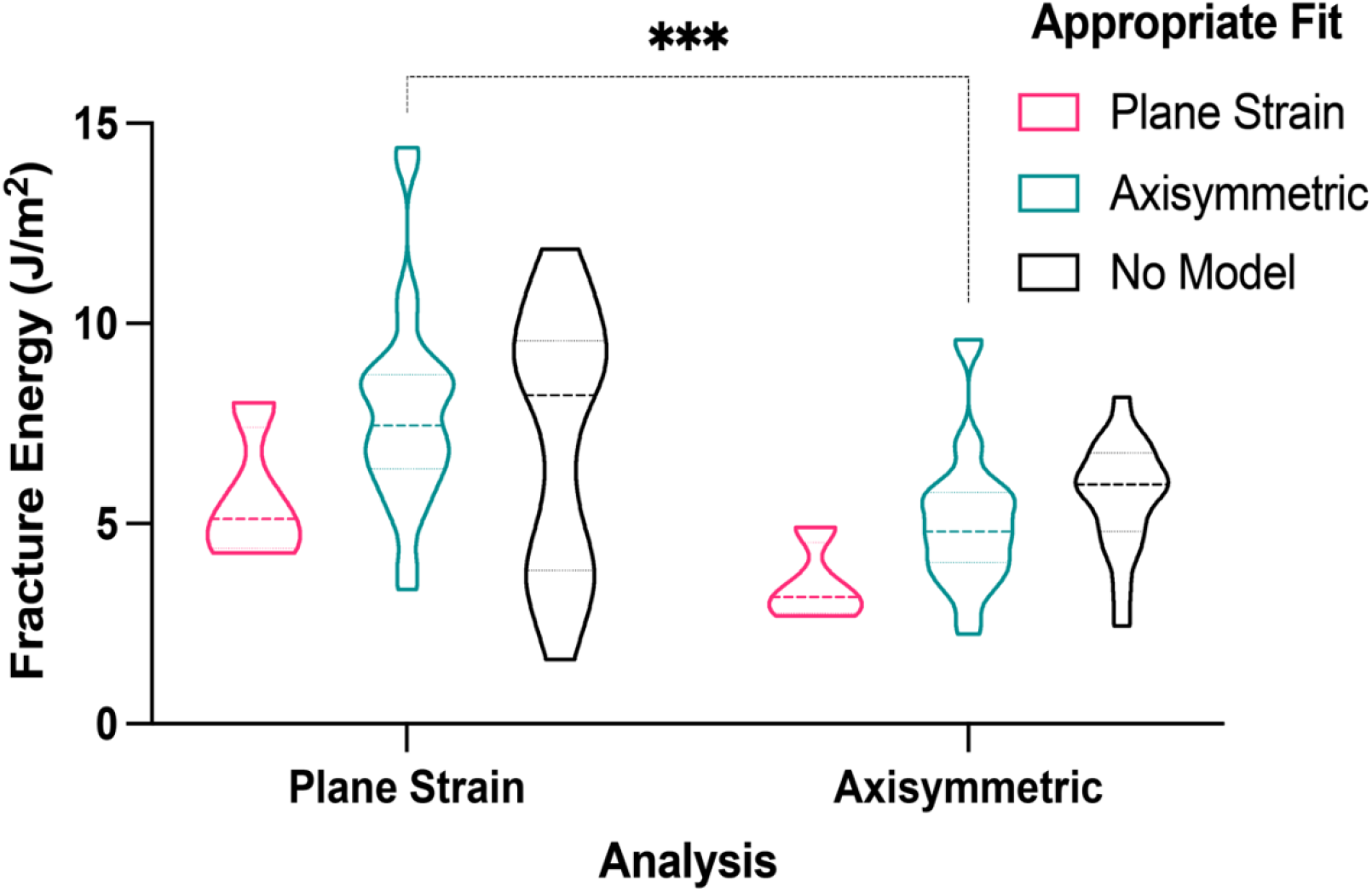
Applying alternate model to each set of data. Applying the plane strain model to experimental data that fits the axisymmetric model results in significantly higher fracture energies, but applying the axisymmetric model to experimental data that fits the plane strain model results in slightly lower values.

### Numerical Simulations of Hydraulic Fracture in Brain Tissue

We developed a finite-element model to simulate the hydraulic fracture of brain tissue using Abaqus (version 2017. Dassault System, Rhode Island). The plane strain model for the KGD case and axisymmetric model for the penny-shaped case are set up in ABAQUS/Standard. Both models have a size of 100 mm by 100 mm. An initial crack size with a length of 0.1 mm is introduced at the edge. To have high accuracy and efficiency, we use extra fine mesh near the initial crack and coarse mesh far from the crack.

In the plane strain model for the KGD case, brain tissue is modeled as a linear poroelastic material with a modulus of 20 kPa, Poisson’s ratio of 0.4, permeability of 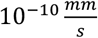, and a void ratio of 0.25 using the CPE4P element. The injection fluid is water with a specific weight of 980 kg/m^3^. The interaction between hydraulic fracture and natural fracture from initiation to propagation was modeled in the cohesive zone model by introducing a layer of cohesive elements (COH2D4P) with a bilinear traction-separation (Figure 4a). In the bilinear constitutive response of traction-separation law, *T* represents the interfacial strength, *δ_c_* is the critical separation, *δ_f_* is the separation at failure, and the area under the curve, *G_c_* is the critical strain energy release rate. In our simulation, a mixed mode fracture is under consideration, but we set the fracture energy of mode II (*G_IIc_*) to ten times larger than the fracture energy of mode I (*G_Ic_*), to only have a mode I fracture. The interfacial strength *T* ranges from 1 to 2 kPa, and the fracture energy of mode I (*G_Ic_*) ranges from 0.0625 to 1 J/m^2^. The initial response of the cohesive element is assumed to be linear until a damage initiation criterion is met. The penalty stiffness *K_i_* of the bilinear traction-separation law is defined as

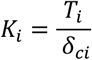

where *i* = 1 stands for tensile deformation and *i* = 2 stands for shear deformation. The definition of *i* is also applied for the following equations. We choose the quadratic stress as the damage initiation criterion:

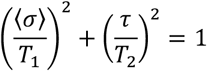

where *σ* is the tensile stress, *τ* is the shear stress, and the Macaulay bracket 〈*σ*〉 represents that the compressive stress does not contribute to the damage initiation. We assume the interfacial strength of shear deformation is the same as the tensile strength. Once the damage initiates, the stiffness begins to degrade. The softening response of the cohesive element is

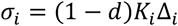

where *d* is scalar stiffness degradation (SDEG in Abaqus), which equals 0 when the interface is undamaged, and *1* when the interface is fully fractured. The energy-based Benzeggagh and Kenane (BK) damage evolution criterion is adopted

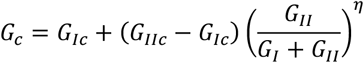

where the BK material parameter *η* is 1.45. In the standard BK option in Abaqus, as the accumulated energy release rate *G*(*G* = *G_I_* + *G_II_*) is larger than the critical energy release rate *G_c_*, the interface is fully fractured. Here, *G_I_* and *G_II_* are the calculated energy release rates for mode I and mode II, respectively.

For the axisymmetric model, brain tissue is modeled using the CAX4P element and the interaction between hydraulic fracture and natural fracture is simulated by cohesive zone model with COHAX4P cohesive element. In the simulations, we use a brain tissue modulus of 20 kPa, an interfacial strength of 0.3 to 2 kPa, and a fracture energy (mode I) of 0.0625 to 1 J/m^2^.

**Supplemental Figure 5.**
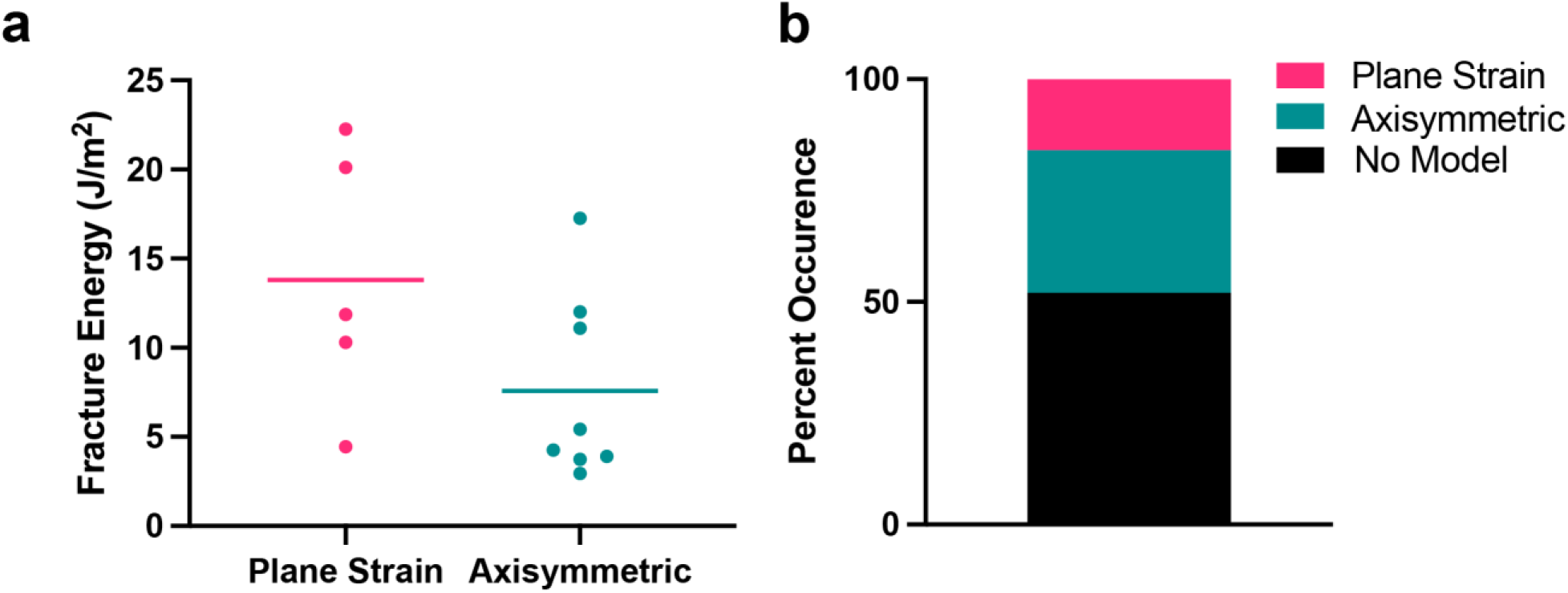
First measurements of lung fracture energy. **a.** The fracture energies of pig lung are consistent regardless of which hydraulic fracture model is applied. **b.** The distribution of the models applied in the pig lung NIC experiments.

**Supplemental Figure 6.**
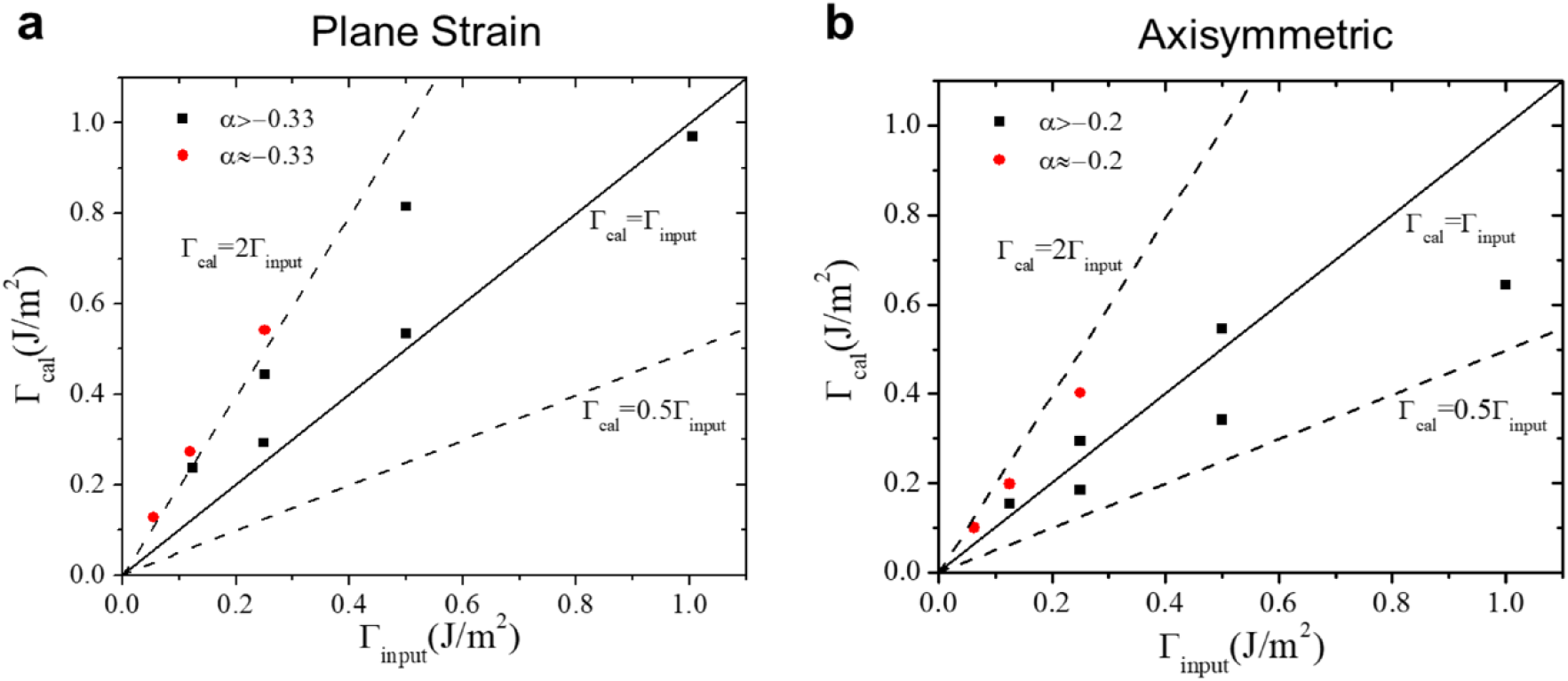
Fracture energy is consistent in simulations and calculations. The calculated fracture energy (Γ_*cal*_) values are within a range of 50% and 200% of the input (Γ_*input*_) values for the **a.** plane strain and **b.** axisymmetric models. Real fracture energy inputs experimental data into simulations and calculated fracture energy uses pressure-time data generated with simulations and the hydraulic fracture equations to calculate fracture energy.

**Supplemental Figure 7.**
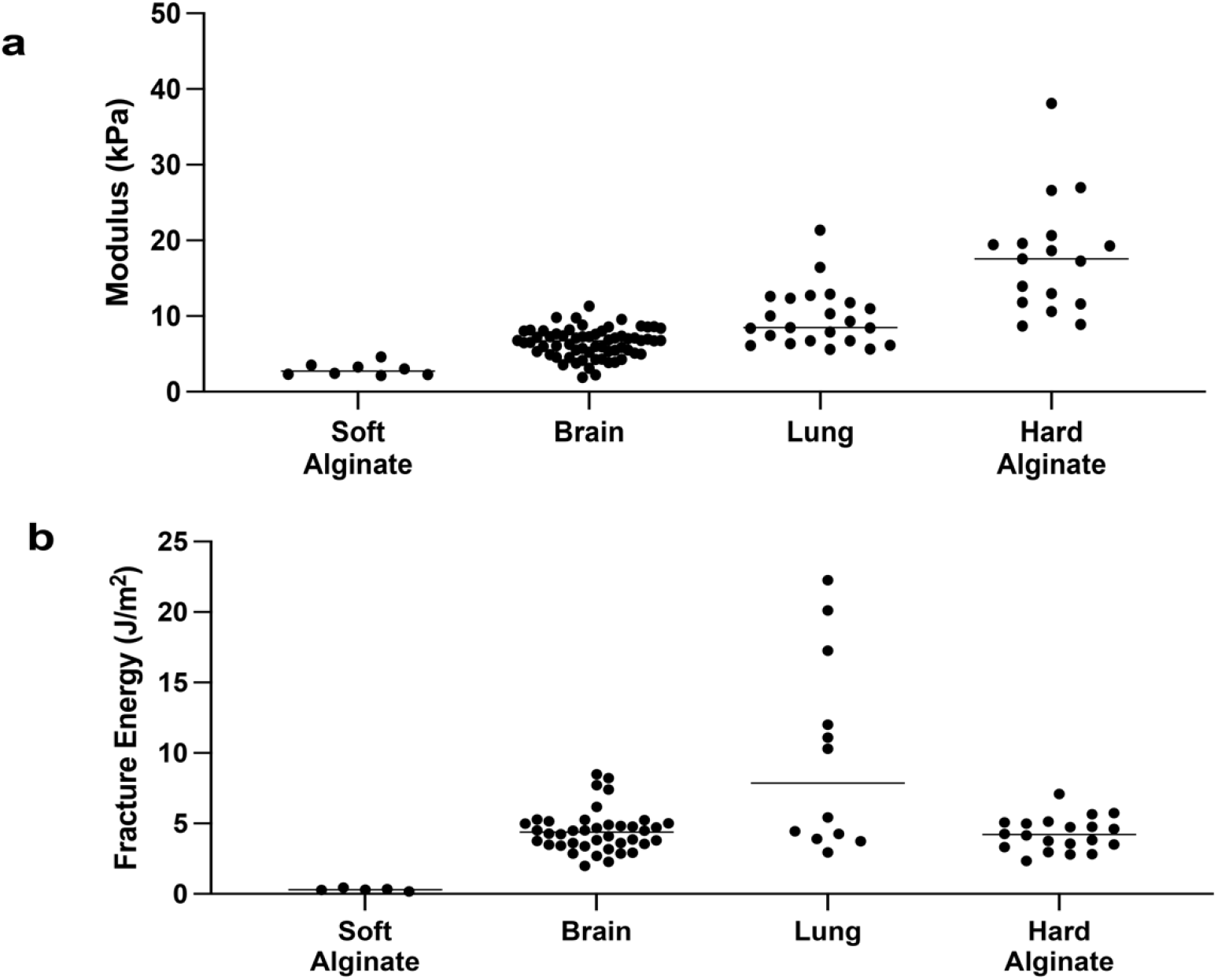
Summary of all modulus and fracture energy values. **a.** NIC measured modulus for soft alginate, brain, lung, and hard alginate do not directly correlate with **b.** fracture energies of the same materials.

#### Gelvatol Mounting Medium Protocol

(Adapted from: https://medschool.ucsd.edu/research/moores/shared-resources/microscopy/Pages/protocols.aspx#pro3)

Materials:
Polyvinyl alcohol
Glycerol
Distilled water
1 L Pyrex beaker
Heat source
Weigh dish

Methods:

1. To a pyrex beaker, add 24 g Polyvinyl alcohol to 60 glycerol and stir
2. Add 60 mL distilled water and leave for several hours to dissolve at room temperature covered
3. Add 120 mL of 0.2 M Tris-Cl (pH 8.5)
4. Heat (to no more than 50°C with occasional stirring for 10 minutes
5. When most of Gelvatol dissolves, clarify by centrifugation at 5000 G for 15 min
6. Aliquot into 15 mL conical tubes, parafilm the tubes to seal, and store at 4°C

**Supplemental Video 1.**
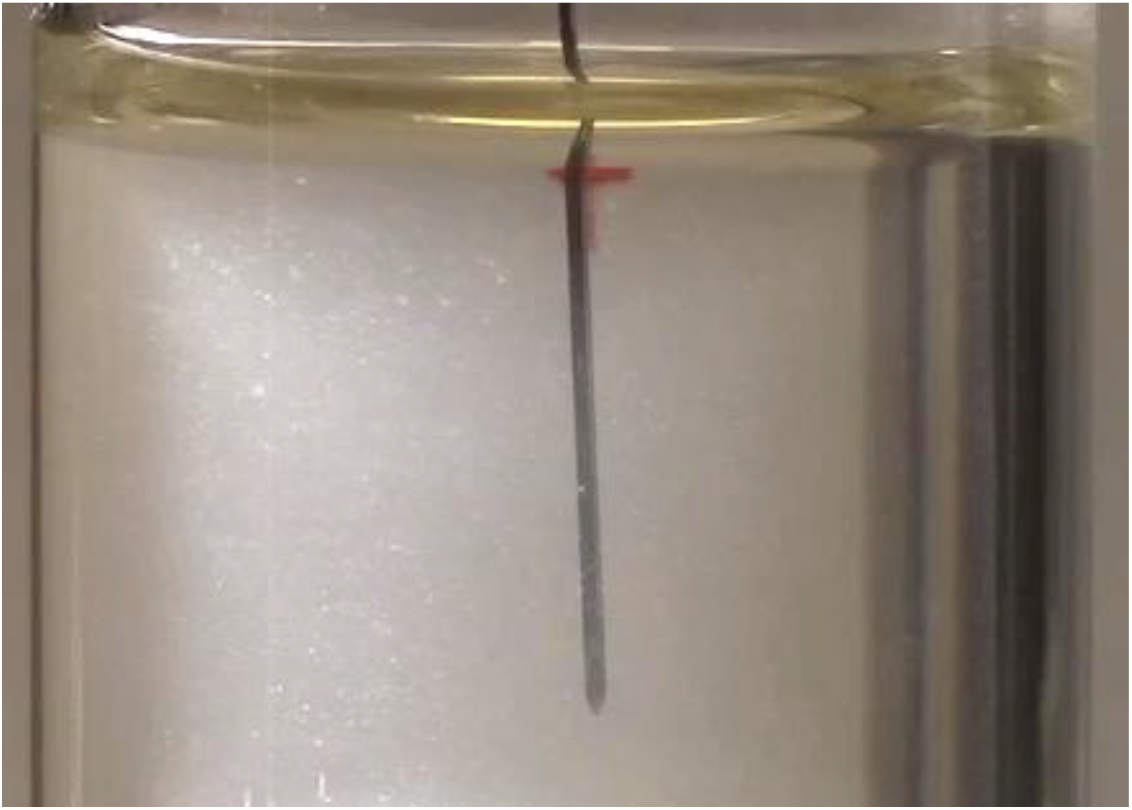
NIC fracture of soft alginate gel.

**Supplemental Video 2.**
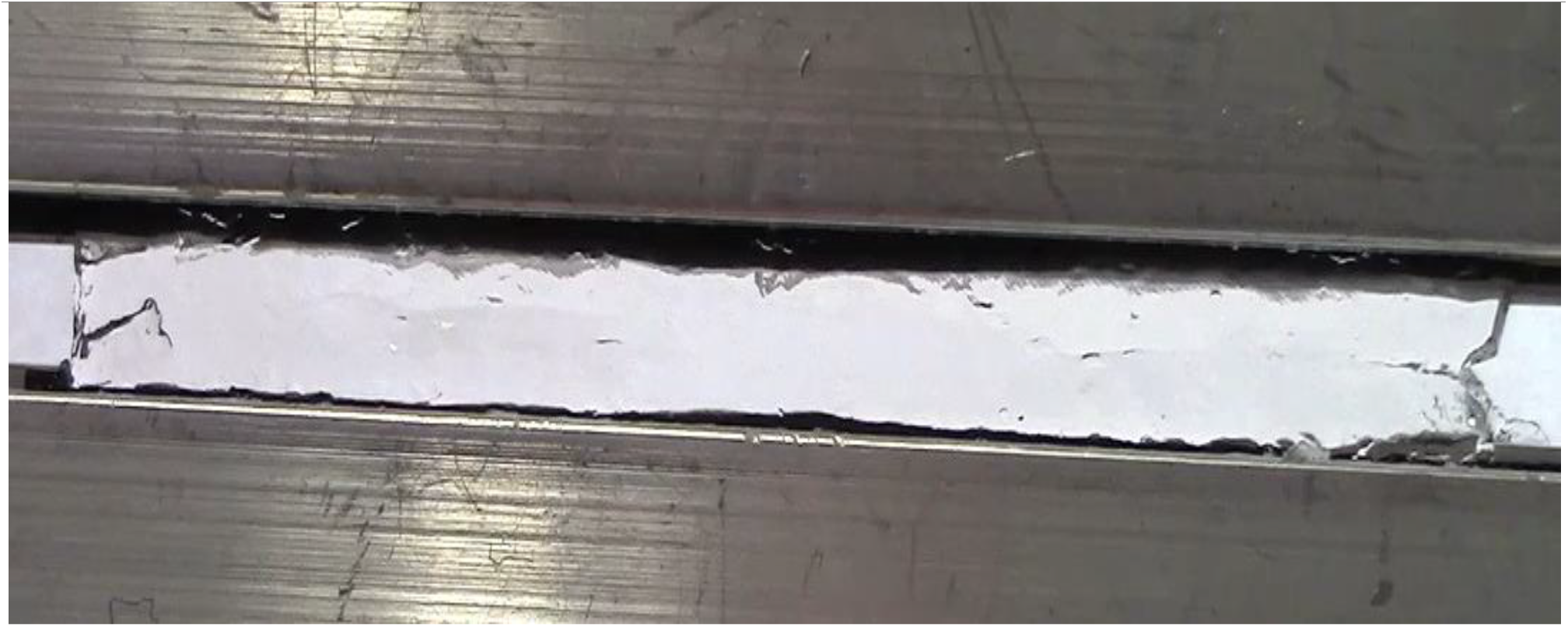
Pure shear of soft alginate gel.

**Supplemental Video 3.**
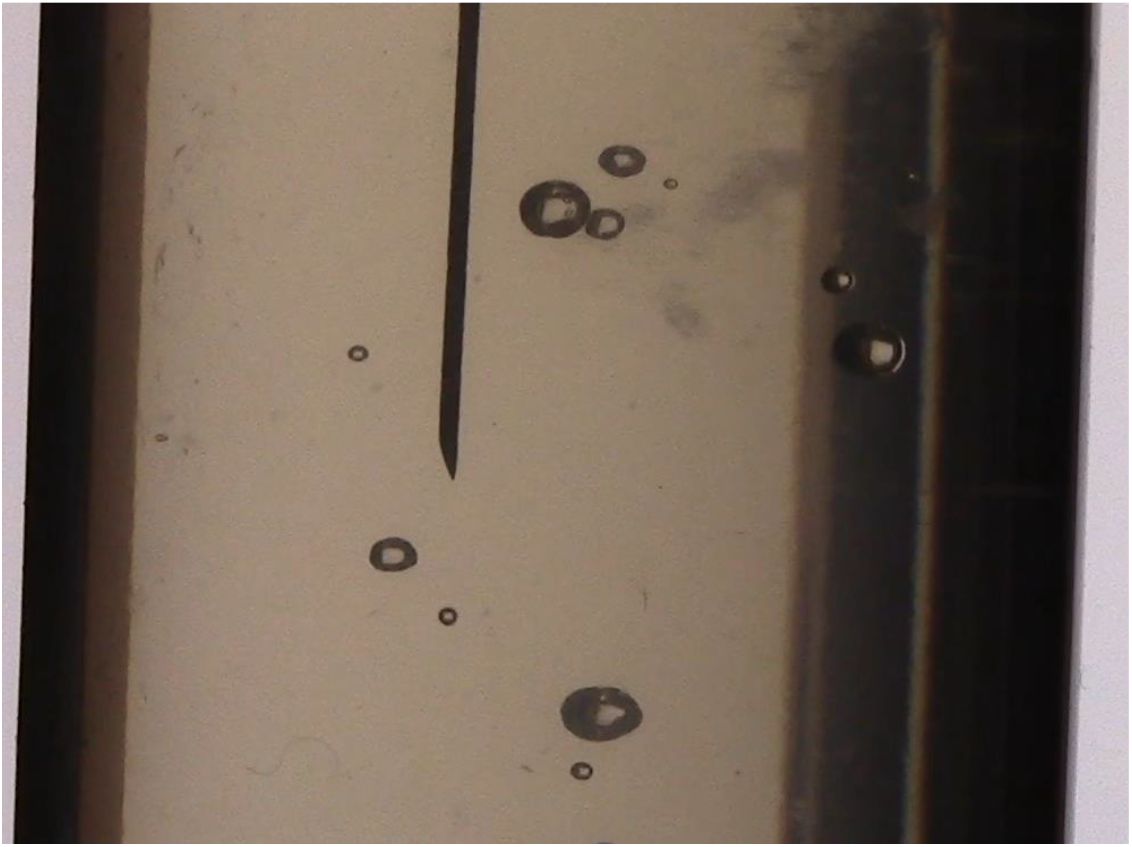
NIC fracture of hard alginate gel.

**Supplemental Video 4.**
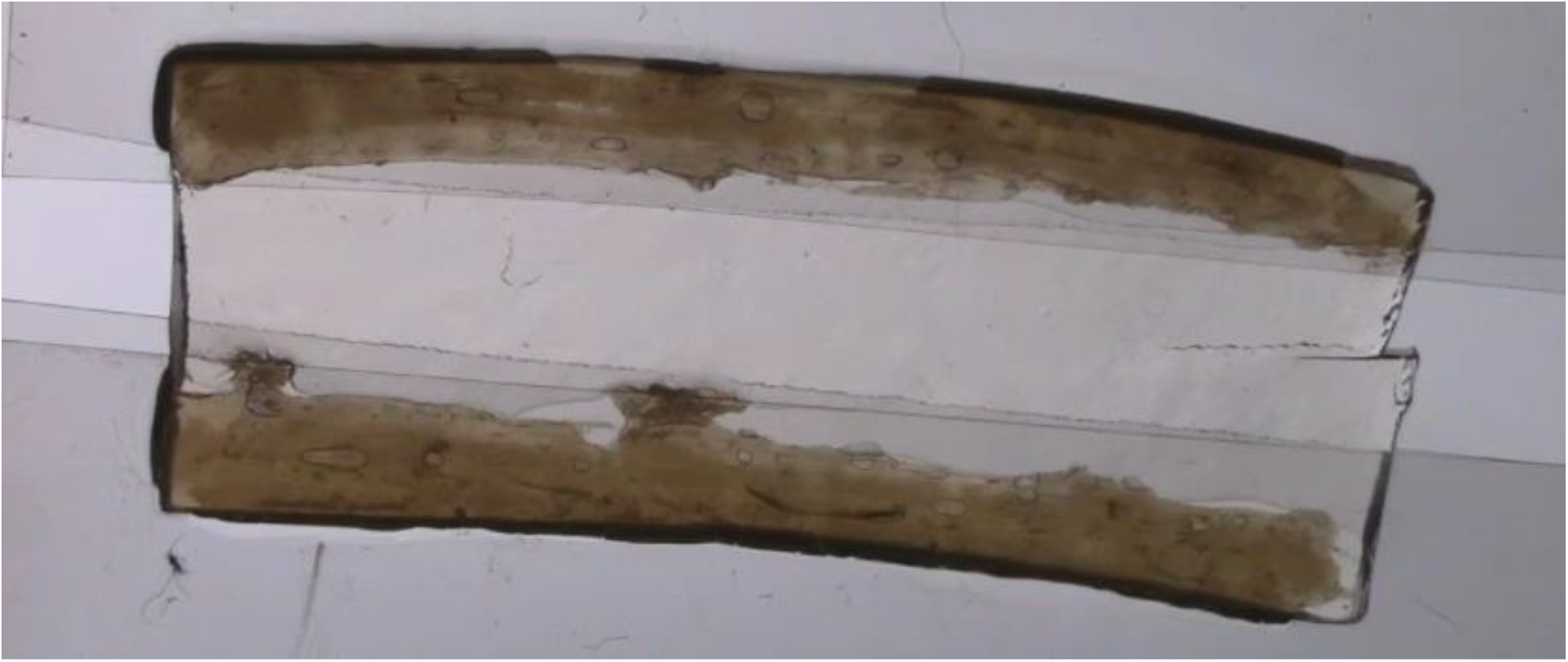
Pure shear of hard alginate gel.

